# Mitogen Kinase Kinase (MKK7) controls cytokine production *in vitro* and *in vivo* in mice

**DOI:** 10.1101/2021.06.25.449967

**Authors:** Amada D. Caliz, Hyung-Jin Yoo, Anastassiia Vertii, Cathy Tournier, Roger J. Davis, John F. Keaney, Shashi Kant

## Abstract

Mitogen kinase kinase 4 (MKK4) and Mitogen kinase kinase 7 (MKK7) are members of the MAP2K family which can activate downstream mitogen-activated protein kinases (MAPKs). MKK4 has been implicated in the activation of both, c-Jun N-terminal Kinase (JNK) and p38 MAPK, whereas MKK7 only activates JNK in response to different stimuli. The stimuli as well as cell type determine the choice of MAP2K member that mediates the response. In a variety of cell types, the MKK7 contributes to the activation of downstream MAPKs, JNK, which is known to regulate essential cellular processes, such as cell death, differentiation, stress response, and cytokine secretion. Previous studies have implicated the role of MKK7 in stress signaling pathways and cytokine production. However, little is known about the degree to which MKK7 and MKK4 contributes to innate immune response in macrophages as well as during inflammation *in vivo*. To address this question and elucidate the role of MKK7 and MKK4 in macrophage and *in vivo*, we developed MKK7- and MKK4-deficient mouse models with tamoxifen-inducible Rosa26 Cre^ERT^. This study reports that MKK7 is required for JNK activation both *in vitro* and *in vivo*. Additionally, we demonstrated that MKK7 in macrophages is necessary for LPS induced cytokine production and migration which appears to be a major contributor to the inflammatory response *in vivo*. Whereas MKK4 plays a significant but minor role in cytokine production *in vivo*.

## Introduction

The mitogen-activated protein kinases (MAPKs) are an evolutionarily conserved signaling mechanism that controls a range of important cellular functions, including differentiation, stress response, and apoptosis [1, 2]. Stress-activated MAPK is comprised of two major pathways, c-Jun N-terminal Kinase (JNK) and p38 MAP Kinase (p38) [3]. JNK and p38 are well-studied signaling cascades that are essential for vital cellular activities [4]. Several members of the mitogen-activated protein kinases kinase (MAP2K) tier of kinases were shown to activate both the JNK and p38 pathways [5–7]. Such diversity in the repertoire of upstream activators might, in turn, define the specificity of the stimuli-induced pathway activation. Mitogen kinase kinase 4 (MKK4) and MKK7, both members of the MAP2K family, MKK4 have been implicated in the activation of downstream stress-activated MAPK JNK and p38 [3, 4, 8, 9]. MKK7 mainly activates downstream kinase JNK in the presence of different stimuli [3, 4]. The cell type and activation stimuli determine which MAP2K, MKK7 or MKK4, plays a major role in the downstream targets of JNK and p38 activation *in vitro* and *in vivo* [10].

The innate immune response is an evolutionarily conserved response and the first line of defense against harmful pathogens [11]. The inflammatory response is a well-orchestrated event initiated by pathogens, such as bacteria, and mediated by a quick activation of MAPK kinases which leads to the production of cytokines as well as the amplification of an inflammatory response affecting both the immune and non-immune cells [12]. Understanding the cell-type- and stimuli-specific mechanisms responsible for the signal transduction is essential for managing the uncontrolled inflammatory responses. The precise contribution of MKK4 and MKK7 in the macrophage-specific immune response has not been discerned. Using both *in vivo* and *in vitro* models, we determined the contribution of each kinase with respect to downstream target activation, cytokine secretion, and macrophage migration; all essential features of an effective immune response.

Pro-inflammatory cytokines, such as tumor necrosis factor-alpha (TNFα) and bacterial lipopolysaccharides (LPS), activate MAPK, JNK and p38 [13]. However, it is not clear which MAP2K plays a larger role during TNFα and LPS activation of JNK and p38 in macrophages. The purpose of this study is to determine whether MKK4 or MKK7 plays a critical role in JNK activation and cytokine production during inflammation in macrophages. In this study we have shown that MKK7 plays a major role in TNFα and LPS-activated JNK signaling and cytokine production *in vitro* and *in vivo*. Although MKK4 contributes to LPS inflammatory response in macrophages, its effects on the JNK signaling and cytokine production are modest compared to MKK7. Data from this study indicates that the MKK7 pathway represents a potential target for the development of therapeutic drugs that may be beneficial for the treatment of sepsis and inflammation.

## Result

### Conditional MKK7 generation

JNK kinases can be activated by different MAP2Ks, which include MKK4 and MKK7 proteins [2, 14]. p38 MAPK can be activated mainly by MKK3 and MKK6 but MKK4 can also plays role in some circumstances [8]. But role of MKK7 in activation of p38 MAPK is not known. Previously, we have established the role of mixed lineage kinases (MLKs) mitogen-activated protein kinases kinase kinase (MAP3K) in TNFα and LPS-activated JNK and p38 MAPKs *in vitro* and *in vivo* [4, 13, 15]. MLKs also regulate cytokine production *in vitro* and *in vivo*. Additionally, activation of JNK and p38 via MKK4 and MKK7 has been implicated in cytokine production [10]. Nonetheless, the roles of MKK4 and MKK7 in LPS-induced inflammation *in vivo* and in macrophages are not known. To investigate the role of MKK4 and MKK7 in LPS-induced cytokine production *in vivo*, we used our previously published MKK4 conditional mice [16] and generated MKK7 conditional mice (Suppl Fig. 1). More specifically the temporal deletion in MKK4 and MKK7 mice was achieved by crossing MKK4-floxed and MKK7-floxed mice with inducible Rosa26-Cre^ERT^ mice (Suppl Fig. 1). Genotypic analysis utilizing different tissues isolated from tamoxifen-treated Rosa26-Cre^ERT^-*Mkk4*-floxed (*Mkk4*^Δ/Δ^) and Rosa26-Cre^ERT^ - *Mkk7*-floxed (*Mkk7*^Δ/Δ^) mice, demonstrated the specific disruption of the *Mkk4* and *Mkk7* genes (Suppl Fig. 1) which was not observed in control mice. This finding was confirmed by immunoblot analysis of MKK4 and MKK7 expression in bone marrow drive macrophages (BMDMs) isolated from control or *Mkk4*^Δ/Δ^ and *Mkk7*^Δ/Δ^ mice treated with 4-hydroxytamoxifen (Suppl Fig. 1).

### MKK7 is required for TNFα-stimulated MAP kinase activation

A pro-inflammatory cytokine TNFα binds to TNFα receptors (TNFR) on the cell surface and activates signal transduction mechanisms leading to phosphorylation of MKK4 and MKK7 protein kinases [10]. We used mouse embryonic primary fibroblast cells (MEFs) treated with TNFα to examine the contribution of MKK4 and MKK7 to p38 and JNK MAPK activation in macrophages. First, we isolated mouse embryonic fibroblast (MEFs) from control or Rosa26-Cre^ERT^-*Mkk4*-floxed and Rosa26-Cre^ERT^-*Mkk7*-floxed mice. These MEFs were treated with 4-hydroxytamoxifen for specific gene deletion. Using these MKK4 and MKK7 knockout MEFs, we assessed the contribution of MKK4 and MKK7 into MEFs by stimulating these cells with TNFα and compared them to Rosa26 Cre^ERT^ control cells. We found that deficiency of MKK4 caused only minor changes in TNFα-stimulated MAP kinase activation JNK (Fig. 1A). Conversely, TNFα-stimulated MAP kinase activation of JNK in *Mkk7*^Δ/Δ^ MEFs was markedly reduced (Fig. 1A). Next, bone marrow-derived primary macrophages (BMDMs) isolated from Rosa26-Cre^ERT^-*Mkk4*-floxed and Rosa26-Cre^ERT^-*Mkk7*-floxed conditional mice were treated with the 4-hydroxytamoxifen for respective gene deletion. Similar to the MEFs, treatment with TNFα to MKK4-deficient BMDMs caused only minor changes in TNFα-stimulated JNK activation (Fig. 1B). However, TNFα-stimulated JNK activation was severely affected by MKK7 deficiency compared to the controls (Fig. 1B). We also observed a minor change of activation in p38 MAPK in MKK7-deficient BMDMs. Together, these data indicates that the majority of TNFα-stimulated MAP kinase activation is mediated by MKK7; while MKK4 acts as a minor partner.

**Figure 1.**
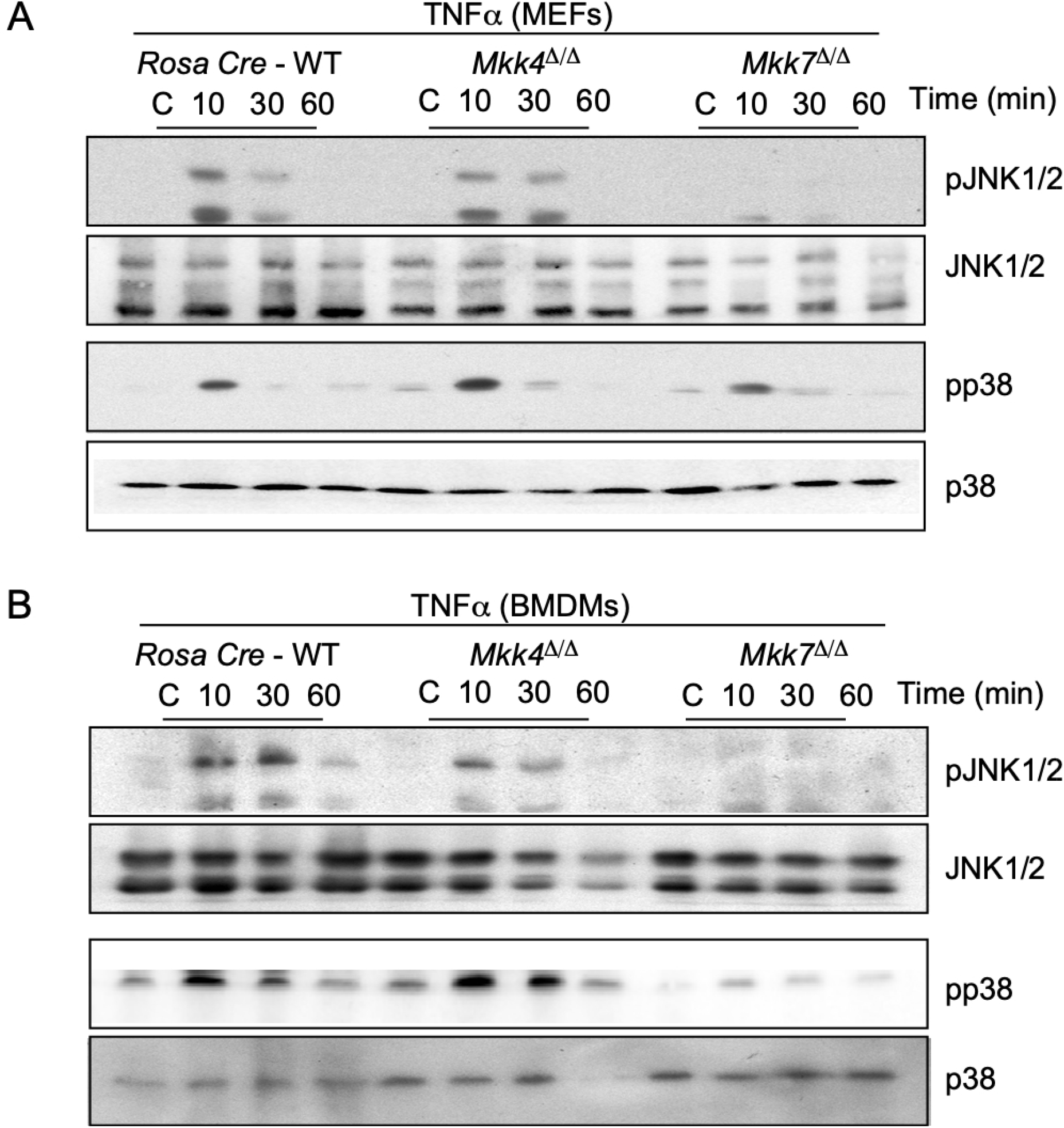
TNF causes activation of a MKK7 signal transduction pathway. (A) MEFs isolated from Rosa26 -Cre^ERT^, Rosa26-Cre^ERT^-*Mkk4*-floxed and Rosa26-Cre^ERT^-*Mkk7*-floxed conditional mice were treated with the 4-hydroxytamoxifen for respective gene deletion. After 5 days of deletion MEFs were treated with or without 10 ng/mL TNFα across three timepoints. MAP kinase JNK and p38 activation were examined by immunoblot analysis. (B) BMDMs from different genotypes were treated with or without 10 ng/mL TNFα across three timepoints (in minutes). JNK and p38 MAPK activation were examined by immunoblot analysis.

### MKK7 plays an important role in LPS-induced MAP kinase activation and cytokine production

To further examine the individual contributions of MKK4 and MKK7 in LPS-induced MAPK activation in macrophages, we compared the responses of JNK and p38 activation in wild-type control, *Mkk4*^Δ/Δ^ and *Mkk7*^Δ/Δ^ BMDMs to the LPS treatment. Our data showed that similar to TNFα-induced MEFs, MKK4 deficiency produces only a moderate decrease in JNK and p38 activation in LPS-activated macrophages (Fig. 2). In contrast, MKK7 deficiency leads to a significant reduction in both JNK and p38 activation (Fig. 2).

**Figure 2.**
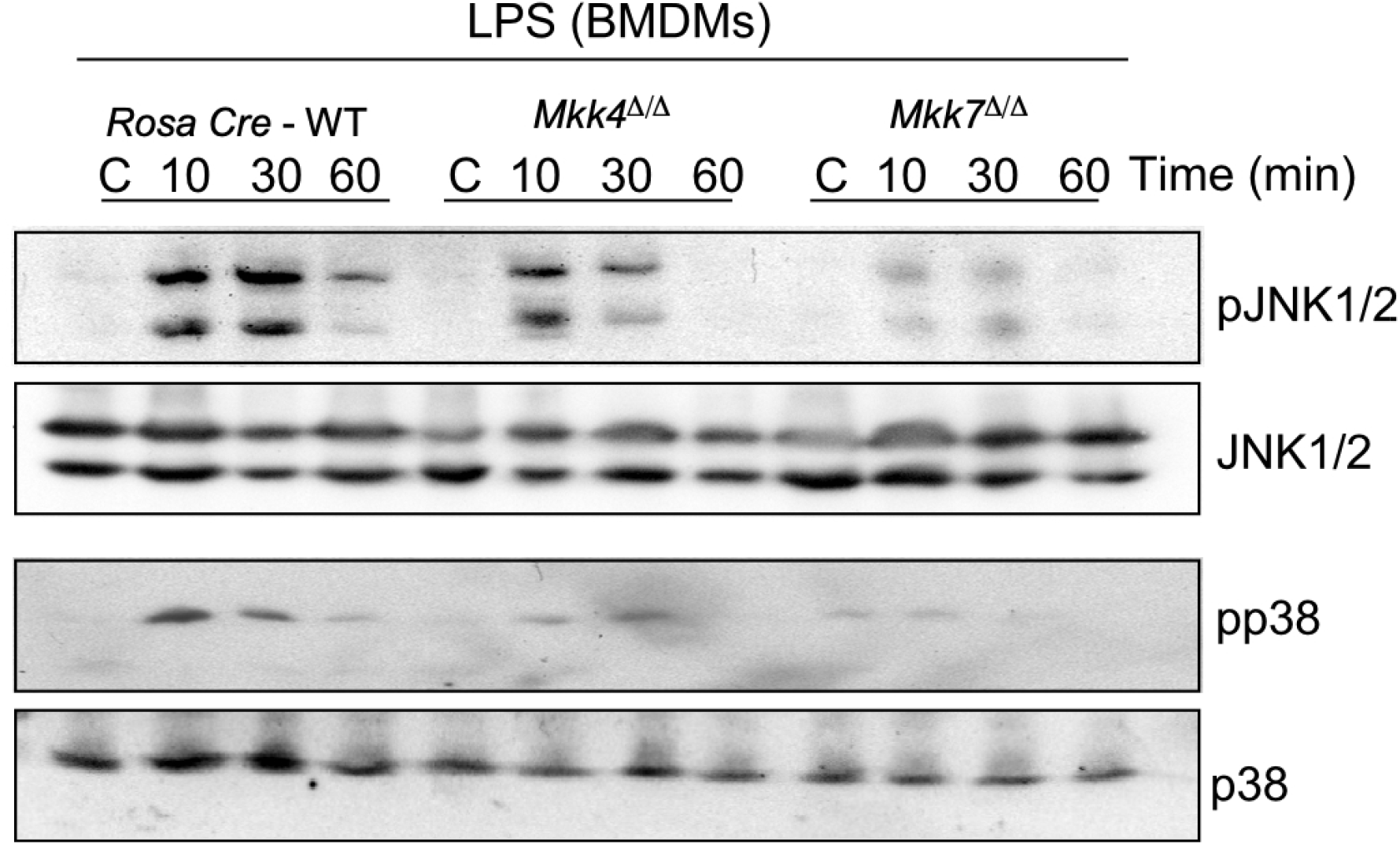
Effect of MKK4 or MKK7 -deficiency on the response to LPS. BMDMs isolated from Rosa26 -Cre^ERT^, Rosa26-Cre^ERT^-*Mkk4*-floxed and Rosa26-Cre^ERT^-*Mkk7*-floxed conditional mice were treated with the 4-hydroxytamoxifen for respective gene deletion. After 5 days of deletion BMDMs were treated without or with 100 ng/ml LPS for 10, 30 and 60 minutes. Protein extracts were examined by immunoblot analysis by probing with antibodies to MAP kinases and phospho-MAP kinases.

Next, we compared the production of inflammatory cytokines during LPS response in control, MKK4, and MKK7 -deficient BMDMs. We found that both LPS-treated MKK4 and MKK7-deficient BMDMs secrete markedly less TNFα and interleukin 6 (IL6) than the wild-type control cells (Fig. 3A). However, the reduction in cytokine production was significantly higher in *Mkk7*^Δ/Δ^ than *Mkk4*^Δ/Δ^ (Fig 3A). Next, we analyzed mRNA production of TNFα and IL6 pro-inflammatory cytokines in both MKK4 and MKK7-deficient BMDMs. Like the secreted cytokines proteins, mRNA expression was also compromised in both MKK4 and MKK7-deficient BMDMs when compared to control cells. Consistent with cytokine secretion, the effect of MKK7-deficiency caused a greater decrease in TNFα and IL6 cytokine expression than MKK4 -deficiency in BMDMs (Fig. 3B). Collectively, these data show that MKK4 and MKK7 are required for cytokine production in LPS induced macrophages with MKK7 having the stronger impact on pro-inflammatory cytokines at the mRNA and protein levels.

**Figure 3.**
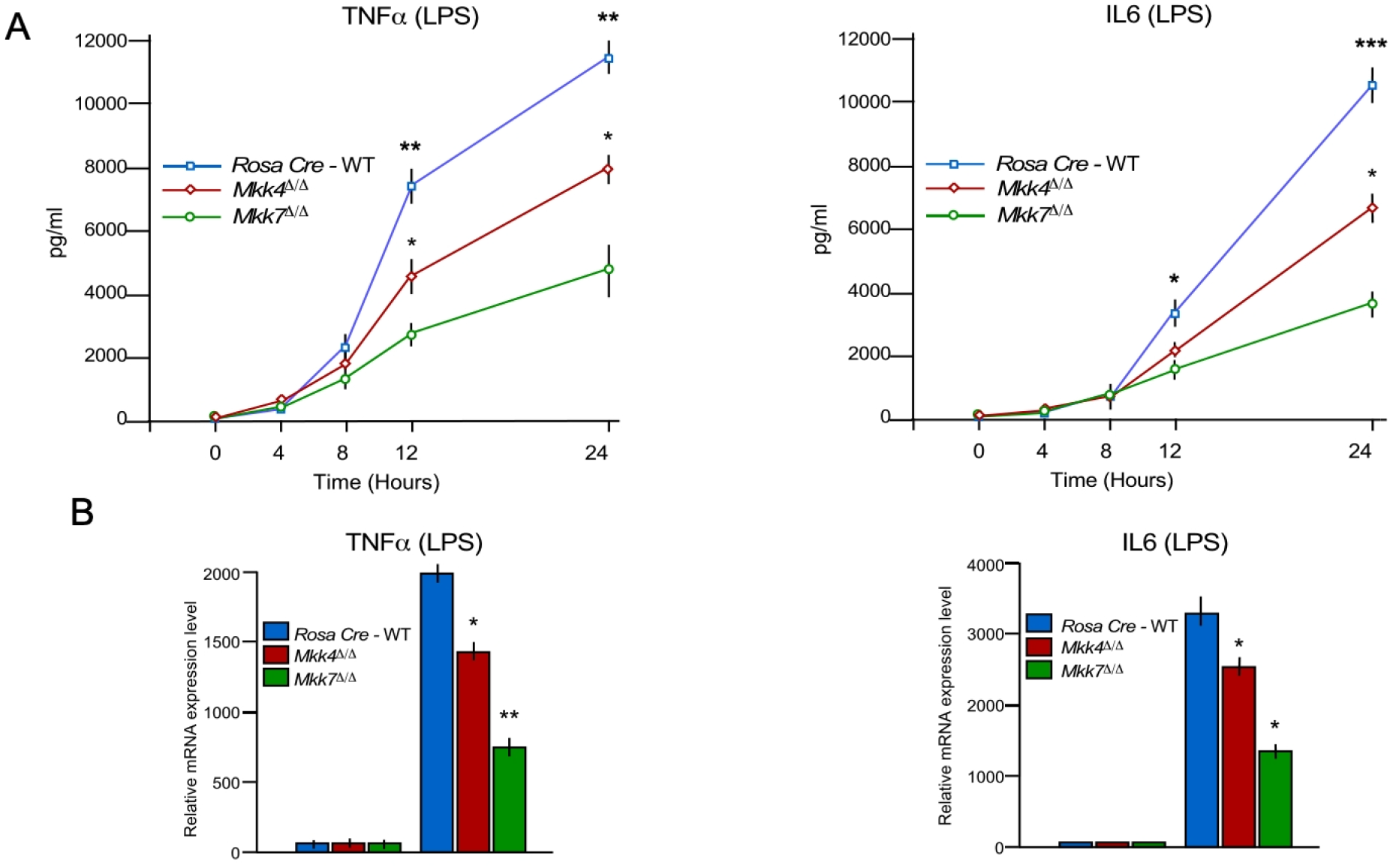
The MKK4 and MKK7 pathway contributes to LPS-stimulated and cytokine production in macrophages. (A-B) Rosa26 -Cre^ERT^, *Mkk4*^Δ/Δ^ and *Mkk7*^Δ/Δ^ BMDMs were treated with 100 ng/mL LPS. The amount of TNFα and IL6 in the culture medium was measured by ELISA (A) and RT-qPCR was performed to measure mRNA production of TNFα and IL6 (B) (SE; n = 6). Statistically significant differences between groups are indicated. (*) P < 0.05; (**) P < 0.01; (***) P < 0.001.

### MKK 7 controls cell migration

MAPK signaling has been implicated in macrophage migration and invasion. Healthy macrophages require robust migration and invasion to execute their proper function; thus, highlighting the importance of determining the role of MKK4 and MKK7 in macrophage migration and invasion. We found that MKK4 deficiency leads to a modest disruption of cell migration and invasion in macrophages (Fig. 4A) while MKK7-deficiency produced a major blockage in the migration and invasion of macrophages compared to the control (Fig. 4A). These data demonstrate that MKK7 is a key component of macrophage migratory machinery. Next, we checked the role of the MKK7 downstream target, JNK, during the migration and invasion of macrophages. We used 4-hydroxytamoxifen treated Rosa26-Cre^ERT^-*Jnk1*-floxed *Jnk2*-/- mice (*Jnk1*^Δ/Δ^ *Jnk2*-/-) for these experiments. Like MKK7 -deficiency, JNK1/2 -deficiency in macrophages leads to a significant reduction of macrophage migration and invasion (Fig. 4B).

**Figure 4.**
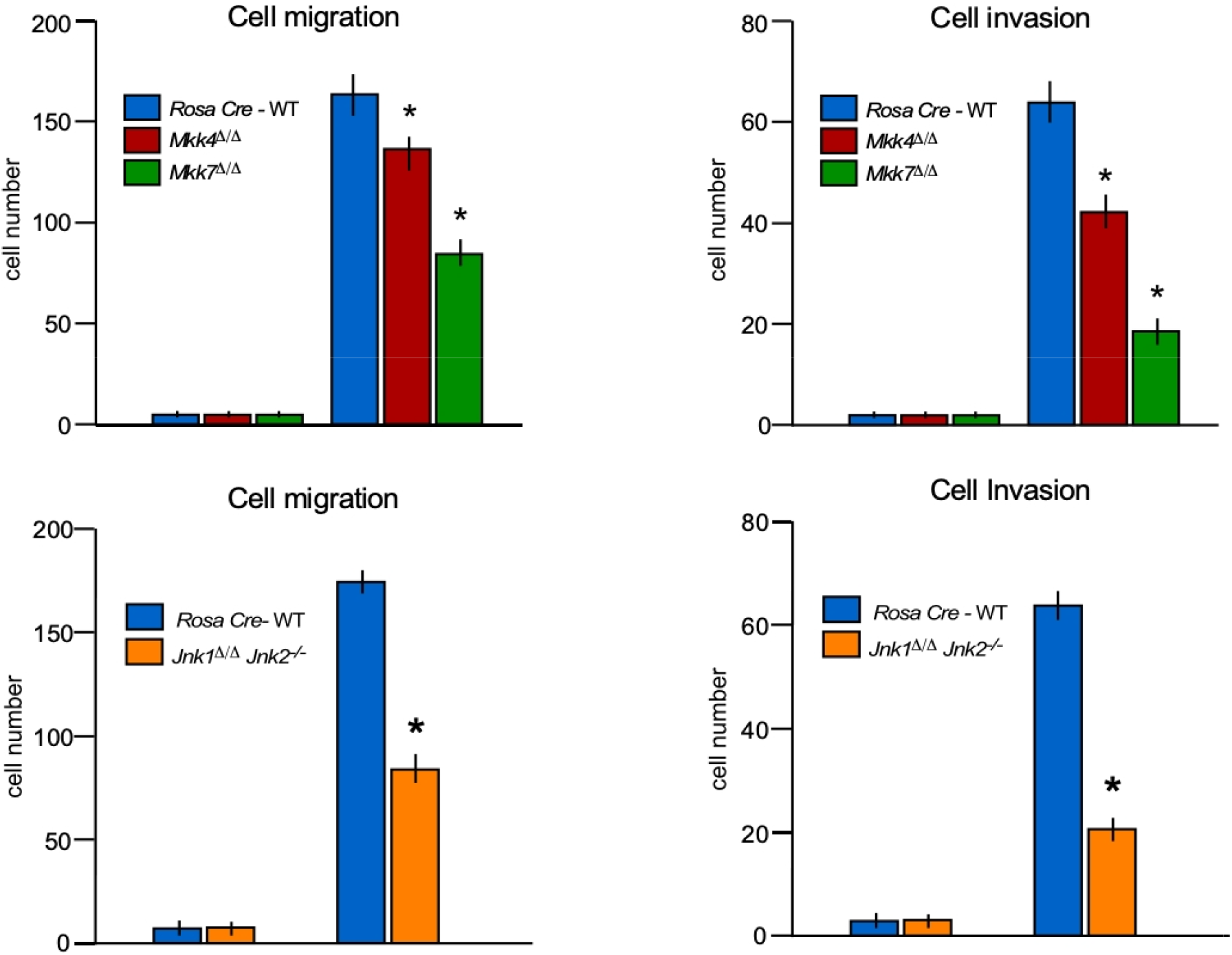
Role of MKK4 and MKK7 in cell migration and invasion. (A) Transwell system was used to measure the migration and Matrigel invasion assay of Rosa26 -Cre^ERT^, *Mkk4*^Δ/Δ^ and *Mkk7*^Δ/Δ^ BMDMs. (B) Rosa26 -Cre^ERT^, *Jnk1*^Δ/Δ^ *Jnk2*-/- BMDMs were used to measure the migration and Matrigel invasion assay by Transwell system. (SE; n = 8); (*) p < 0.05.

### MKK7 contributes to inflammatory cytokine production *in vivo*

To examine the contribution of MKK4 and MKK7 to the immune response *in vivo*, we tested the effects of MKK4 and MKK7 deficiency on the response of mice to endotoxin exposure. Rosa26-Cre^ERT^-*Mkk4*-floxed and Rosa26-Cre^ERT^-*Mkk7*-floxed conditional mice were injected with the single dose of tamoxifen for respective gene deletion. Treatment of wild-type mice with LPS caused increased expression of inflammatory cytokines TNFα, IL6, IL1α, and IL1β in the blood (Fig. 5). The production of LPS-induce pro-inflammatory cytokines *in vivo*, specifically, TNFα, IL6, IL1α, and IL1β was significantly suppressed in both *Mkk4*^Δ/Δ^ and *Mkk7*^Δ/Δ^ mice (<u>Fig. 5</u>). Although the decreases in cytokine production in both *Mkk4*^Δ/Δ^ and *Mkk7*^Δ/Δ^ mice were significantly lower compared to the control, the decrease in TNFα, IL6, IL1α, and IL1β production in MKK4 -deficient BMDMs were less compared to MKK7 -deficient mice (Fig. 5). Together, these data establish the MKK4 and MKK7 pathways as contributors to inflammation, and like isolated macrophages *in vitro*, MKK7 has a significantly stronger contribution to the LPS response than MKK4 *in vivo*.

**Figure 5.**
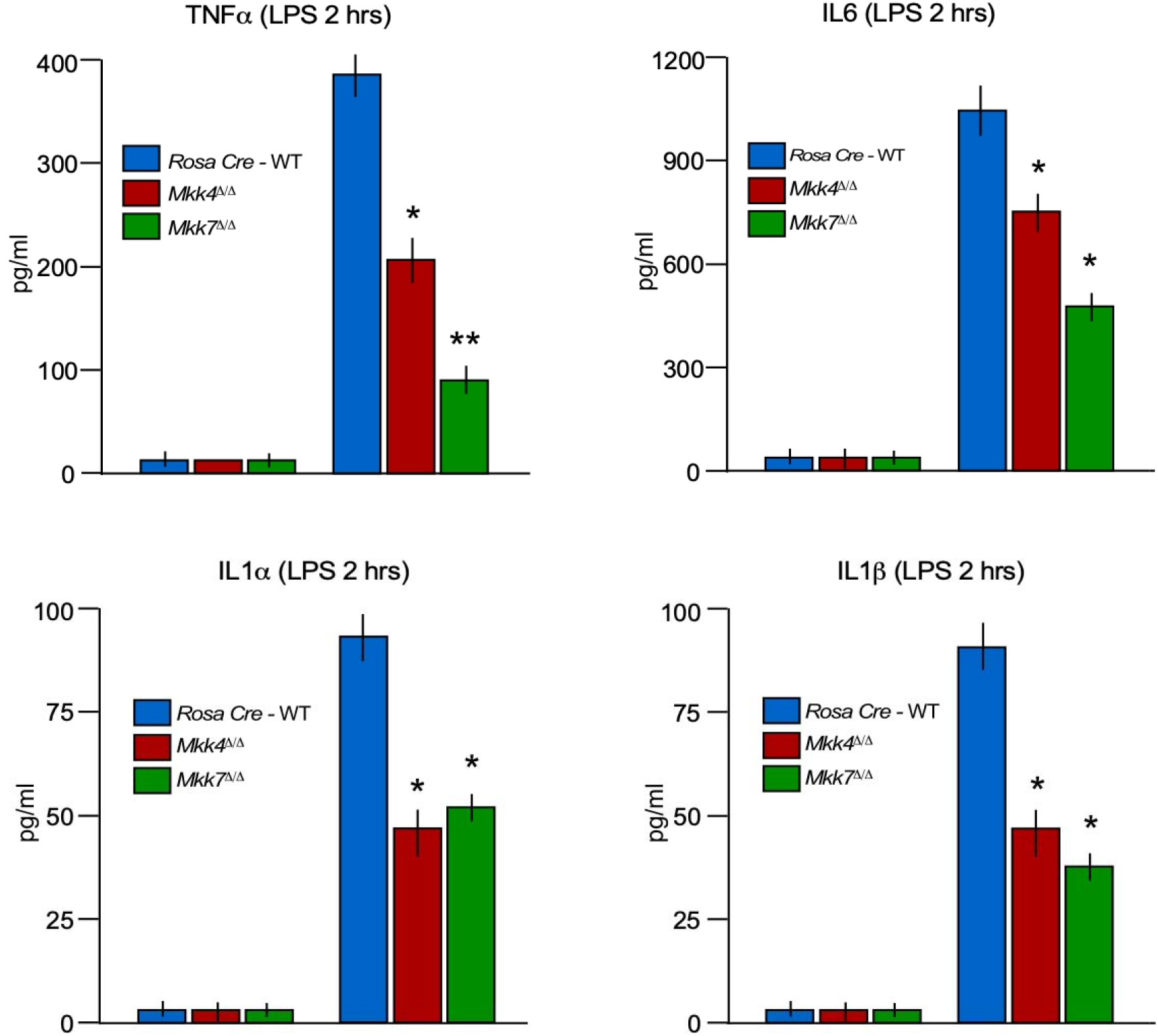
The MKK4 and MKK7 pathway contributes to LPS-mediated inflammation. (A) Rosa26 -Cre^ERT^, *Mkk4*^Δ/Δ^ and *Mkk7*^Δ/Δ^ mice, were treated with or without 20 mg/kg LPS by intraperitoneal injection (2 h). The amount of TNFα, IL6, IL1α and IL1β in the serum was measured by ELISA (SE; n = 8). Statistically significant differences between wild-type and MLK-deficient mice are indicated. (*) P < 0.05; (**) P < 0.01.

### JNK1/2 mimics the MKK7 during inflammatory cytokine production *in vitro* and *in vivo*

JNK signaling is severely affected by MKK7-deficiency (Fig 1). To dissect the role of JNK1/2 in macrophages, we isolated bone marrow-derived primary macrophages (BMDMs) from Rosa26-Cre^ERT^-*Jnk1/2*-floxed *Jnk2*-/- mice and treated them with 4-hydroxytamoxifen for gene deletion (*Jnk1*^Δ/Δ^ *Jnk2*-/-). Similar to MKK7, LPS-treated JNK1/2-deficient BMDMs secrete markedly less TNFα and IL6 than wild-type cells (Fig. 6).

**Figure 6.**
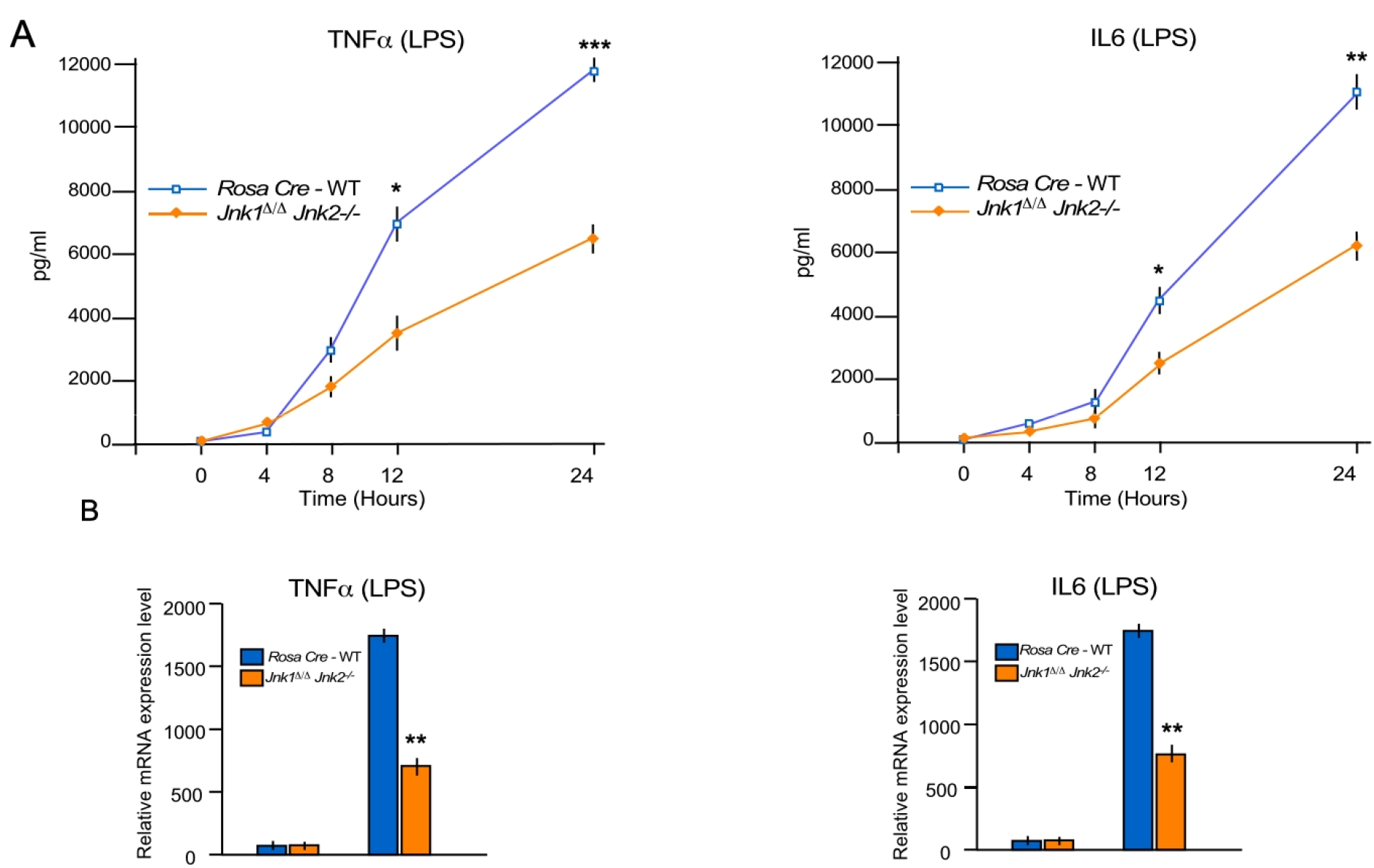
The JNK1/2 mimic the MKK7 phenotype and contributes to LPS-stimulated and cytokine production in macrophages. (A-B) Rosa26 -Cre^ERT^ and *Jnk1*^Δ/Δ^ *Jnk2*-/- BMDMs were treated with 100 ng/mL LPS. The amount of TNFα and IL6 in the culture medium was measured by ELISA (A) and RT-qPCR was performed to measure mRNA production of TNFα and IL6 (B) (SE; n = 6). Statistically significant differences between groups are indicated. (*) P < 0.05; (**) P < 0.01; (***) P < 0.001.

Next, we checked the mRNA expression of the pro-inflammatory cytokines TNFα and IL6 in JNK1/2-deficient BMDMs and found that similar to the secreted cytokines, mRNA expression was reduced in JNK1/2-deficient BMDMs compared to the control cells (Fig. 6). These data suggest that JNK1/2 is required for cytokine production in LPS-induced macrophages.

## Discussion

The MKK4 and MKK7 pathways are mediators of cytokine signaling which results in JNK and p38 MAPK activation. Furthermore, leading to the transcriptional activation of cytokine genes and eventually cytokine production [3, 10]. Biochemical studies using cultured primary cells have provided substantial evidence to support this conclusion [5, 10] as mice lacking *mkk4* or *mkk7* die [5, 7, 10, 17] before birth. Therefore, there has been limited research investigating the roles of MKK4 and MKK7 *in vivo*. To overcome this problem, we have generated conditional MKK4 and MKK7 knockout in this study to examine the roles and contributions of the MKK4 and MKK7 pathways in macrophage activation as well as cytokine production *in vitro* and *in vivo*.

Dissecting the cell type-specific mechanisms that dominate the innate immune response is critical for the successful management strategies of the uncontrolled inflammatory response [11, 18]. Notably, our *in vitro* and *in vivo* data showed that MKK7 plays a major role in comparison to a minor role played by MKK4 during LPS-induced cytokine production. Our data also demonstrated that MKK7 regulates cell migration, an essential feature of efficient macrophage functionality. We also observed that JNK mimics the LPS-induced cytokine phenotype caused by MKK7 deficiency.

MAPK and NFκB pathways play an essential role in cytokine production [19]. Specifically, both innate immune and adaptive cells such as macrophages and T-cells produce various pro-inflammatory cytokines in a MAPK-dependent manner during infection, causing an inflammatory response.

JNK needs phosphorylation on both Thr180 and Tyr182 for its complete activation [5]. JNK phosphorylation occurs preferentially on Tyr182 by MKK4 and on Thr180 by MKK7 [10, 20]. Also, it is known that both MKK4 and MKK7 are required for optimal activation of JNK in fibroblast [5, 10]. MKK4 and MKK7 appear to contribute equally to JNK activation in response to UV radiation [5]. However, MKK7 is essential for JNK activation by TNFα and MKK4 is only required for maximum JNK activation in MEFs [10]. This cooperation of two kinases between MKK7 and MKK4 may be true in macrophages as well in which MKK7 plays a major role in JNK activation and MKK4 is only required to maximize the JNK activation. The role of MKK7 in activation of p38 MAPK has not been established as our data showed that MKK7 might play a minor role in p38 MAPK activation as well. We can speculate that similar to JNK activation, p38 MAPK also requires the assistance of their main activator MKK3/6 [9] with MKK7 for their optimal activation in macrophages.

JNK and p38 are long-studied stress signaling cascades essential for a broad range of vital cellular and organismal activities. Therefore, the use of small molecule inhibitors against JNK or p38 to prevent cytokine production and inflammation would likely be accompanied by severe side effects, making them poor drug targets. Several members of the MAP2K and MAP3K tier of kinases were shown to activate JNK and p38 pathways [4, 13, 21]. Such diversity in the repertoire of upstream activators might, in turn, define the specificity of the stimuli-related pathway activation. Therefore, the data presented in this manuscript suggests that MKK7 plays a vital role in the activation of JNK and TNFα. Moreover, LPS induced cytokine production could lead to the development of drug-targeted treatments for sepsis and inflammation.

## Methods

### Animals

Inducible Rosa26 Cre strain mice were obtained from The Jackson Laboratory (004847). Mice with floxed *Mkk4* gene [16], floxed *Mkk7* (exons 3–6 flanked by LoxP site) [22] and floxed *Jnk1 Jnk2*-/- [23] were bred with the inducible Rosa26-Cre^ERT^ mouse line on the C57BL/6J background (Suppl Fig. 1). Both mice were injected with single dose of tamoxifen (100 mg/kg body weight in corn oil) for respective gene deletion.

All mouse experiments were performed according to the relevant ethical regulations. Mice were housed in a facility accredited by the American Association for Laboratory Animal Care. All animal studies were approved by the Institutional Animal Care and Use Committee of the University of Massachusetts Medical School and the Brigham Women’s Hospital, Boston.

### Cell culture

Bone marrows drive primary macrophages (BMDMs) were prepared and cultured in Dulbecco’s modified Eagle’s medium supplemented with 30% L929 supernatant (source of M-CSF), 20% fetal bovine serum, 100 U/mL penicillin, 100 mg/mL streptomycin, and 2 mM L-glutamine (Invitrogen). Embryonic day 13.5 (E13.5) primary MEFs were isolated and cultured in Dulbecco’s modified Eagle’s medium supplemented with 10% fetal bovine serum, 100 U/mL penicillin, 100 mg/mL streptomycin, and 2 mM L-glutamine (Invitrogen). Both BMDMs and MEFs isolated from different genotyped of mice were treated with 4-hydroxytamoxifen (2uM) for 24 hours for respective gene deletion. Cells were used for experiments after 5 days of 4-hydroxytamoxifen treatment.

### RNA Preparation and Quantitative Polymerase Chain Reaction

Quantitative RT–PCR assays (TaqMan) of mRNA expression were performed using a 7500 Fast Real-Time PCR machine (Applied Biosystems) with total RNA prepared from cells with an RNeasy minikit (Qiagen). TaqMan©assays were used to quantitate *II6* (Mm00446190_m1), *Tnfa* (Mm00443258_m1) and *GAPDH* (4352339E-0904021) mRNA (Applied Biosystems). The 2^-ΔΔ*CT*^ method is used for relative quantification of gene [24, 25]. Reference genes of *GAPDH*, was used to normalize the PCRs in each sample.

### Immunoblot Analysis

Cell extracts were prepared using triton lysis buffer (TLB buffer) [20 mM Tris (pH 7.4), 1% Triton X-100, 10% Glycerol, 137 mM NaCl, 2 mM EDTA, 25 mM β-Glycerophosphate] with proteinase inhibitors (Sigma #11873580001) and phosphatase inhibitors (Sigma #4906837001). Protein extracts (50 μg of protein) in β-mercaptoethanol containing SDS sample buffer were separated in 4% to 12% gradient SDS-polyacrylamide gels (Bio-Rad #456-8094) and transferred to nitrocellulose membranes (Bio-Rad #170-4271, Hercules, CA) and incubated with a primary antibody with 1:1000 dilution. Immunocomplexes were visualized with horseradish peroxidase-conjugated secondary antibodies and detected with a Clarity Western ECL substrate (Bio-Rad #170-5061, Hercules, CA), and images were acquired on a chemiluminescent imager (Bio-Rad Chem-Doc Imaging System).

### Analysis of Blood

Plasma cytokines and cytokines secreted in cell culture media were measured using ELISA (Luminex 200; Millipore) by following the manufacturer’s instructions.

### Antibodies and reagents

Primary antibodies were obtained from Cell Signaling (MKK4, MKK7, phospho-JNK1/2, p38a, phospho-p38, and IκBα), BD Pharmingen (JNK1/2), and Sigma (a-Tubulin, #T5168).

### Statistical Analysis

All data are expressed as mean ± SE, and the numbers of independent experiments are indicated. Statistical comparisons were conducted between 2 groups by use of Student t-test or Mann–Whitney U test as appropriate. Multiple groups were compared with either 1-way Kruskal–Wallis or ANOVA with a post hoc Tukey–Kramer multiple comparisons test as indicated in legends. A P value <0.05 was considered significant. All statistics were done using StatView version 5.0 (SAS Institute, Cary, NC) or GraphPad Prism version 5 (GraphPad Software, La Jolla, CA).

## Data Availability

The data sets generated and analyzed as part of this study are available upon request from the corresponding author.

## Acknowledgments

We thank Jennifer Cederberg, Jason Hagan, Kathy Gemme, and Marisol Diaz for academic assistance, Tammy Barrett for technical assistance. We thank Ana C. Dolan for reading this manuscript. This work was supported by AHA grant 16SDG29660007 (to S.K.) and NIH grants R01 HL151626 (to J.F.K.) and R01 DK107220 (to RJD).

