## Supplemental Figures for "Mitogen Kinase Kinase (MKK7) controls cytokine production *in vitro* and *in vivo* in mice"

**Supplementary Figures**

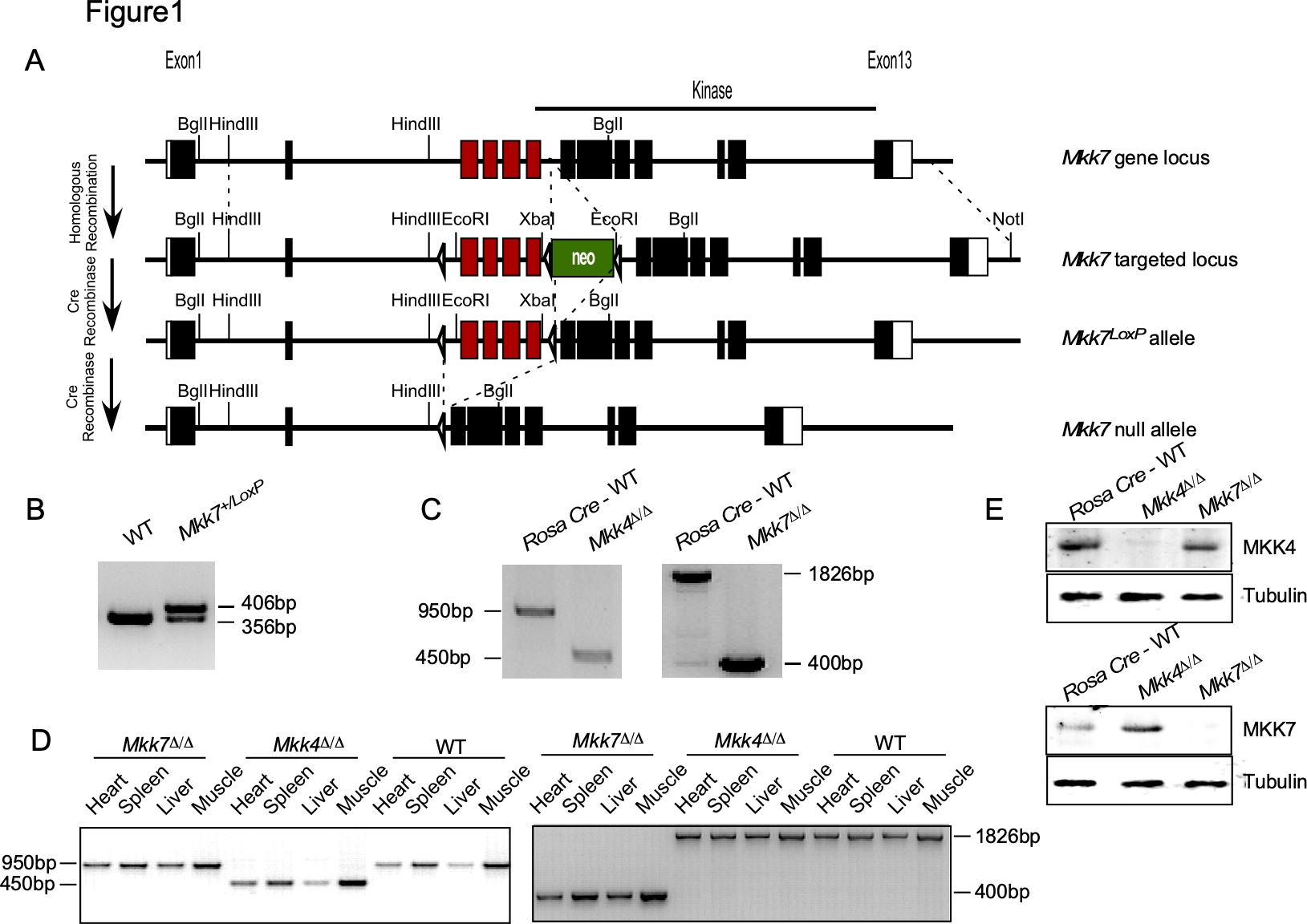

**Supplementary Figure S1. Construction of MKK7-deficient mice.** (A) Strategy for the creation of MKK7-floxed and MKK7 conditional knockout mice. The structure of the MKK7 genomic locus and the targeted locus are illustrated. (B) Genomic DNA isolated from mouse tails was examined by PCR to detect wild-type and Floxed alleles of MKK7 (C-D) Genomic DNA isolated from mouse tails was examined by PCR to detect wild-type and deleted alleles of MKK4 and MKK7 mice (C); Genomic DNA isolated from different tissues were examined by PCR from different genotypes of mice (D). (E) Extracts prepared from Rosa26 -Cre^ERT^, Mkk4^∆/∆^ and Mkk7^∆/∆^ MEFs were examined by immunoblot analysis using antibodies to MKK4, MKK7, and α-Tubulin.
